# Role of finger movement kinematics in friction perception at initial contact with smooth surfaces

**DOI:** 10.1101/2023.10.29.564644

**Authors:** Naqash Afzal, Sophie du Bois de Dunilac, Alastair J. Loutit, Helen O Shea, Pablo Martinez Ulloa, Heba Khamis, Richard M. Vickery, Michaël Wiertlewski, Stephen J. Redmond, Ingvars Birznieks

**Author notes:** **Corresponding author’s name and email address:** Naqash Afzal. **Financial interests or conflicts of interest**: The authors declare no competing financial interests.

## Abstract

When manipulating objects, humans adjust grip force to friction remarkably quickly: it may take just 100 ms to see adjustment to friction at the skin-object interface. While the motor commands adapt, subjects become aware of slipperiness of touched surfaces. In this study, we explore the sensory processes underlying such friction perception when no intentional exploratory sliding movements are present. Previously, we have demonstrated that humans cannot perceive frictional differences when surfaces are brought in contact with an immobilized finger (Khamis et al., 2021b) unless there is a submillimeter lateral displacement (Afzal et al., 2022), or subjects made the movement themselves (Willemet et al., 2021). In the current study, subjects actively interacted with a device that can modulate friction using ultrasound, without an exploratory sliding movement, as they would when gripping an object to lift it. Using a two-alternative forced-choice paradigm, subjects had to indicate which of two surfaces felt more slippery. Subjects could correctly identify the more slippery surface in 87 ± 8% of cases (mean±SD; n=12). Biomechanical analysis of finger pad skin contacting a flat smooth surface indicated that natural movement kinematics (e.g., tangential movement jitter and physiological tremor) may enhance perception of frictional effects. To test whether this is the case, in a second experiment a hand support was introduced to limit fingertip movement deviation from a straight path. Subject performance significantly decreased (66 ± 12% correct, mean±SD; n=12), indicating that friction perception at the initial contact is enhanced or enabled by natural movement kinematics.

**Significance statement:** Sensing surface friction is crucial for automatic grip force control to avoid dropping objects. A slipping handhold can lead to loss of balance and falling. In many instances, the required grip force may exceed hand’s physical ability or an object’s breakage point, therefore cognitive selection of a safe and achievable action plan based on friction perception is critical. Little is known about how our awareness of surface slipperiness is obtained under such circumstances without exploratory movement. The current study demonstrates that natural movement kinematics inducing submillimeter lateral movements play a central enabling role, demonstrating interdependence between the motor system and sensory mechanisms. These findings broaden our fundamental understanding of sensorimotor control and could inform the development of advanced sensor technologies.

## Introduction

A safe grip between the fingertip skin and a surface can only be established if the applied grip force is large enough to create a friction force that is equal to the load force developing tangential to the surface (Johansson and Westling, 1987). The motor control system achieves adequate grip force control by adjusting it to the frictional properties of the surface and skin (Johansson and Flanagan, 2009). When we hold objects like tools in our hands, in addition to automatic motor adjustments, there is also a conscious sensory awareness of how slippery the gripped surface is and whether there is sufficient traction provided by the skin to perform the intended action safely. The sensory mechanisms enabling perception of surface slipperiness under the conditions commonly encountered during object manipulation, without sliding or rubbing movements of the fingertips over the surface, are not well understood.

In previous studies, we investigated whether humans could perceive frictional differences between surfaces just by touching a surface without performing sliding movements (Khamis et al., 2021b). Biomechanical investigations indicate that skin deformation patterns can efficiently reflect the frictional condition (Johansson and Flanagan, 2009; Delhaye et al., 2014; Delhaye et al., 2016; Barrea et al., 2018) and thus could be conveyed by tactile afferent responses (Khamis et al., 2014a; Khamis et al., 2014b). However, when surfaces were brought into contact with an immobilized finger (passive touch), subjects were unable to differentiate the slipperiness of neither smooth nor textured surfaces. We hypothesised that because slipperiness perception in this context is pertinent to manipulation, an active movement might be required to enable sensory perception. Experiments in which subjects themselves actively contacted the surface (active touch) with a finger constrained to move along the normal of the contact surface demonstrated that a radial divergence pattern was sufficient to sense frictional differences; however, this ability depends on optimised contact kinetics and is most efficient when friction is low (Willemet et al., 2021). Further investigations using passive touch, where the object moves to contact the finger, revealed that active movement, in fact, is not necessary to perceive frictional differences, as submillimeter range lateral movement as small as 0.2 - 0.5 mm of the surface relative to the affixed rigid nail-phalangeal bone complex is sufficient to perceive surface slipperiness, even with a larger coefficient of friction (Afzal et al., 2022). Overall, it has become evident that intentional exploratory sliding movements and full slip are not required to perceive surface slipperiness. Variation in outcomes between experimental conditions indicates that the nervous system uses various types of cues and sources of sensory information, depending on availability. For example, the radially divergent skin pattern might be more informative to convey frictional differences with more slippery smooth surfaces when there is little tangential force developing and detection of the resulting lateral slip is difficult (Willemet et al., 2021), while discrimination of surfaces with higher friction, representative of most often-handled objects, may require small lateral movements tangential to the surface.

Natural movements when reaching and gripping objects would rarely occur without the presence of subtle motion jitter which could potentially be the cause of small lateral displacements. We hypothesize that natural movement kinematics producing minor lateral displacements could be the key to enabling us to evaluate the surface slipperiness when gripping and handling an object or tool. In this study, subjects evaluated similar frictional differences, as in our previous study (Khamis et al., 2021b), in which subjects were unable to discriminate between two surfaces when using passive touch, but now asking subjects to touch the surfaces themselves in a natural way. Using fingerprint image processing techniques, we evaluated the presence of partial slip linked to subjects’ ability to discriminate surface slipperiness. To confirm that finger movement kinematics had a decisive role in subjects’ ability to evaluate slipperiness, we performed another experiment in which tangential movement jitter was significantly reduced, and we observed that subjects’ performance also deteriorated significantly.

## Materials and Methods

### Subjects

Twelve healthy right-hand-dominant subjects (mean age 26.1 ± 6.2 years SD, 4 female) participated in the experiment. The experimental protocols were approved by the Human Research Ethics Committee (approval number HC180109) at UNSW Sydney. All the subjects provided written consent before the start of the experiment.

### Friction modulation

An ultrasonic friction reduction device (Wiertlewski et al., 2016) was used to change the friction of a smooth glass surface. The piezoelectric elements of the friction modulation device were driven by a waveform generator (DG 1022; RIGOL Technologies, Beijing, China) via a high-voltage amplifier (A-303; A.A. Lab Systems Ltd., Israel). Three levels of friction high (H), medium (M), and low (L) were obtained by varying the driving voltage amplitude. MATLAB (MathWorks, Natick, MA, USA, 2018a) was used to control the drive voltage amplitude of the friction reduction device via the analog output of a data acquisition unit (USB-6218; National Instruments, TX, USA). The friction reduction device was mounted on a custom-developed platform attached to a six-axis ATI Nano17 force-torque sensor (ATI Industrial Automation, NC, USA). The setup is illustrated in Fig. 1.

**Fig. 1.**
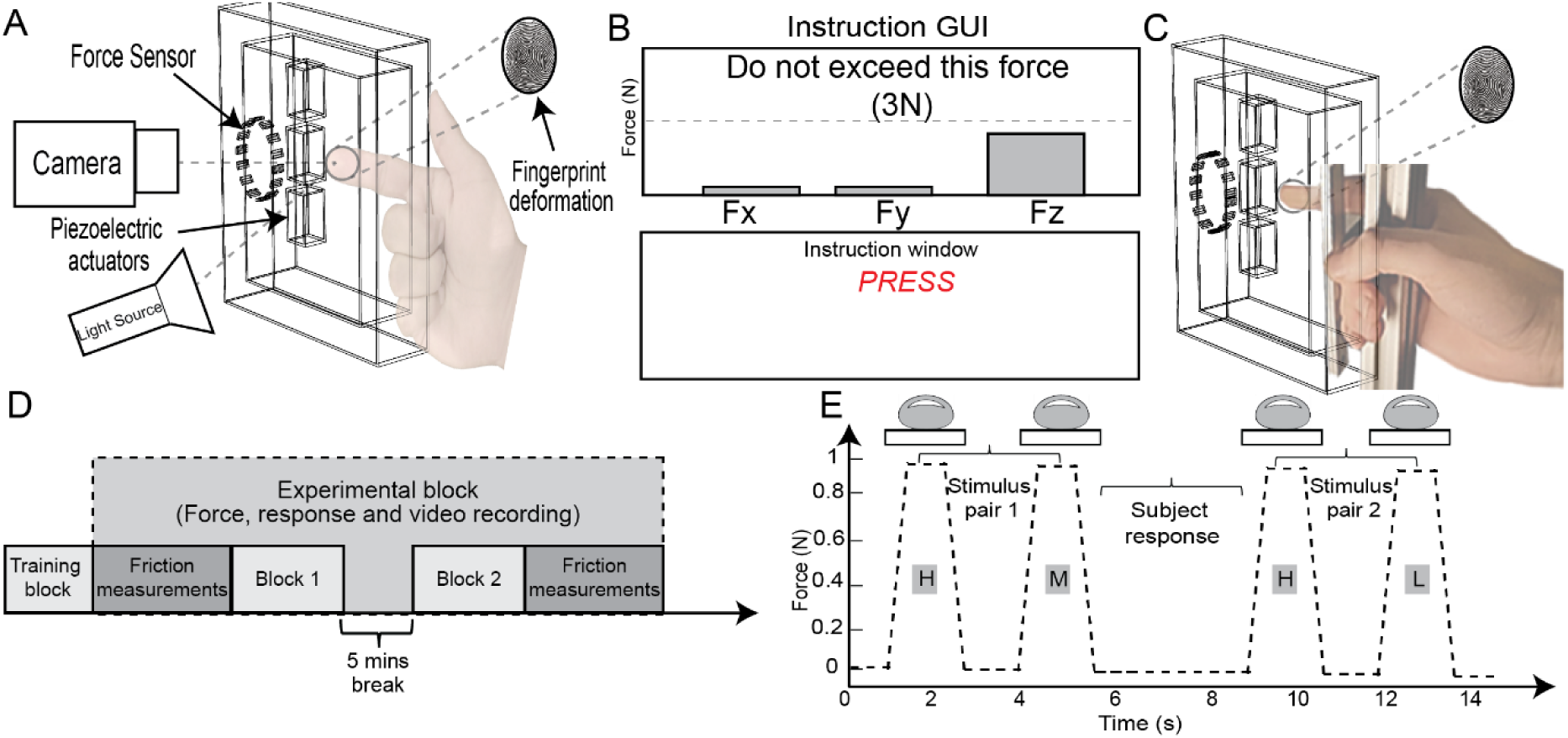
Experimental setup and protocol. *A:* Reach-and-touch condition: illustration of the finger contacting the friction reduction device attached with the piezoelectric actuators modulating the friction of the borosilicate glass mounted on a six-axis force-torque sensor. The light source is positioned under a steep angle to the camera axis to obtain fingerprint images with contrasting ridges and valleys: the light is reflected where there is no skin in contact with the glass of the friction plate, but the light is transmitted (and absorbed by the skin) where skin is in contact with the glass. *B:* A GUI to help pace the timing of when subjects touch, hold, and retract skin from the glass. Display of force levels was shown in real-time bar charts to help subjects maintain the desired force level. *C:* Supported touch condition: Hand holding a vertical t-slot frame with the thumb, palm, and three flexed fingers while touching the friction reduction device using the index finger. *D:* Schematic of the experimental sequence. *E:* Schematic illustration of the time course of presentation and evaluation of the stimulus pairs by subjects touching the friction modulation device (H vs M and H vs L denote pairs of stimuli where H is high friction; L is low friction; M is medium friction). The friction reduction device is switched off for the H condition.

### Data acquisition

Signals from the force/torque sensor, the amplitude of the friction modulation device, and the subject responses were sampled at 1 kHz via another data acquisition unit (PowerLab 16/35; ADInstruments, Australia). The fingertip skin deformation profile within the contact area was analysed using video recorded from a Sony α6300 camera (Sony Corporation, Japan) mounted on a frame.

### Video capture

Videos of the area of contact between the finger pad and the friction plate were acquired at either 60 frames per second (FPS) (subjects 1 to 5) or 120 FPS (subjects 6 to 12). The camera had a resolution of 1920 × 1080 pixels and was mounted in a way that resulted in one pixel per 40×40 µm^2^ of the scene. High contrast between fingerprint ridges and valleys was obtained by utilizing an optical setup that utilized the total internal reflection principle (Fig. 1A) (Tada and Kanade, 2004).

### Experimental procedure

The subjects were seated comfortably in a height-adjustable chair. The subjects’ task was to touch without sliding the vertically-oriented surface of the ultrasonic friction reduction device using the index finger of their right (dominant) hand (Fig. 1A). A two-alternative forced-choice protocol (2AFC) was used for the psychophysics study, wherein two stimuli in a pair, each with a different level of friction, were generated. The timing of when to touch the surface and retract was communicated to subjects by commands on a computer monitor controlled by a GUI developed in MATLAB. The force target was shown on the computer monitor using force magnitude progress bars as shown in Fig. 1B. The trial started with a command of “Press” with a beep sound. The subject was then expected to touch the friction reduction device with a target force of about 1 N. The target force was conveyed to the subjects through the force progress bars on the GUI. A beep sound was generated if the contact force exceeded the 3 N, which happened very rarely. One second after the contact with the surface of the glass was detected, a command to “Lift” the finger was given in conjunction with the sound cue (beep) for retraction of the finger. After retracting the finger, the next stimulus with a different friction level was presented after a 1-second interval. After the presentation of both stimuli in the pair, subjects verbally indicated which of the two stimuli they felt was more slippery. The responses were recorded by the experimenter.

A training block, comprising ten pairs of stimuli, was conducted before the start of the experiment to familiarize the subjects with the task. Two different conditions were tested with the same protocol: (a) Reach-and-touch condition; (b) Supported-touch condition. In the reach-and-touch condition, subjects were asked to actively touch the ultrasonic friction reduction device using the right index finger without any exploratory sliding movements. The complete 2AFC protocol was tested where subjects were instructed to reach and touch the surface of the friction reduction device (Fig. 1A). In the supported-touch condition, a hand support was introduced to minimize any tangential movements relative to the surface typically present during a reach and touch movement due to small trajectory deviations from a straight path towards the surface and physiological tremor originating from arm muscles. Subjects were instructed to hold a vertical aluminium t-slot frame with their thumb, palm, and three flexed fingers while touching the ultrasonic friction reduction device using their index finger (Fig. 1C). The same 2AFC protocol was used to test subjects’ ability to perceive frictional differences.

Each condition was tested with a total of 60 stimulus pairs. The 60 stimulus pairs for each condition were divided into two experimental blocks, each block comprising 30 stimulus pairs with a five-minute break between each block. Each pair of stimuli was presented twenty times; in ten trials the higher friction was presented first followed by the lower friction, and in the other ten trials the lower friction was presented first followed by the higher friction, all presented in a pseudorandom order. Subjects were asked to wear headphones playing white noise to mask auditory cues from the equipment; however, the instructional beeps had sufficient volume to be heard above this white noise. The schematic of the whole experimental protocol is shown in Fig. 1D.

### Friction measurements and detection of frictional differences by sliding movement

The friction between the fingertip and the contact surface depends upon the accumulation of sweat and several other factors including skin properties (hydration, elasticity, smoothness, etc.). Ten measurements at each friction level (H, M, and L) were obtained before and after each experimental block. The stimulus presentation procedure and subject’s task were similar to the rest of the trials, except that at the end of each trial instead of lifting their finger off, the subjects were instructed to slide the finger inwards (proximal) to generate a slip. We determined the static coefficient of friction (µ_s_) by measuring the tangential-to-normal force ratio at the time when the whole fingertip contact area slipped. The time of slip was determined by visual inspection of force traces and video recording of the fingerprint at the moment when the load force began decreasing.

### Image processing

Fingerprint images (Fig. 2A) extracted from the video recording were used to detect local movement of the skin over the glass plate of the friction modulation device. Video frames were corrected for perspective by tracking the position of four screws fixed with respect to the glass plate, and a rectangular region of interest encompassing the contact area during the whole contact duration was cropped out. For each touch, a reference video frame was defined as the frame in which the normal force applied was maximal during the contact with the surface. From that reference time, each video recording was split in two sections, before and after the reference frame. Video frames from the reference to the release of the contact, when the finger was retracted, were processed forward in time, and frames from first contact (0.1 N normal force) to the reference frame were processed backward; this is done so as to initiate the tracking algorithm at a time when the grip force was largest and the most fingerprint image features are expected to be present and well identifiable for tracking until they disappear as the region of contact between the skin and the glass shrinks. To enhance contrast in the contact area, the grayscale of each frame was adjusted to cover the range of values present in the contact area of the reference frame. The region of contact between the finger pad and the glass plate was then segmented using a combination of Otsu’s thresholding method followed by mathematical morphological operations. The region in which to search for the contact area in a frame was limited to a dilated version of the previous contact area. Within the contact area, features as defined by the minimum eigenvalue algorithm (Shi and Tomasi, 1994) were tracked using the Kanade-Lucas-Tomasi algorithm (Lucas and Kanade, 1981; Tomasi and Kanade, 1991). If the tracking of a feature was interrupted, and resulting trajectories lasted fewer than five frames, those trajectories were discarded from further analysis.

**Fig. 2.**
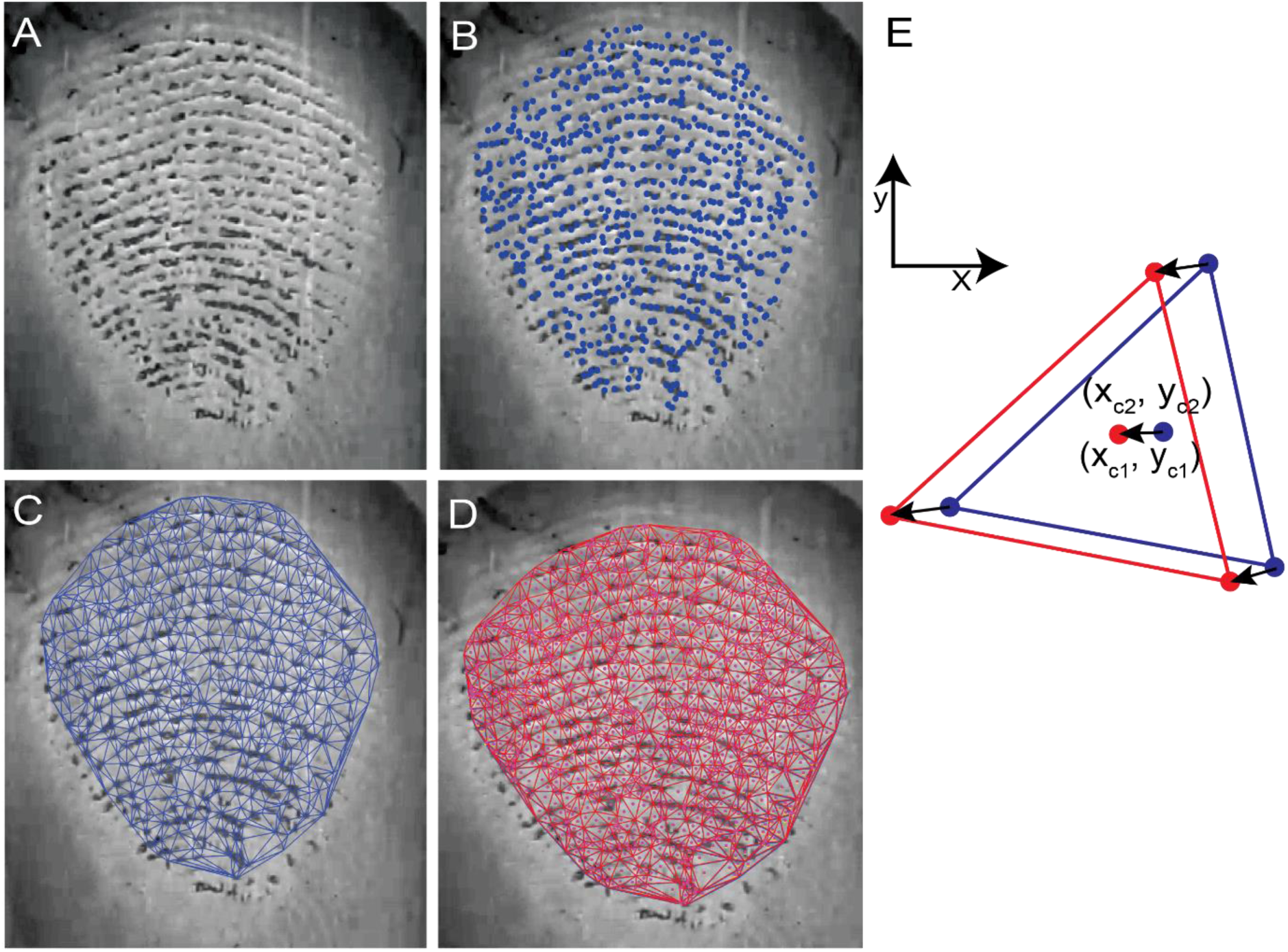
Determining displacement of the skin by fingerprint image analyses. A: An image of the fingerprint within the contact area. B: Identified feature points used for tracking are shown in blue superimposed on the fingerprint image. C: Delaunay triangles are constructed at the current frame joining the tracked feature points (blue triangles). D: Delaunay triangles are constructed for the same features which were detected in the next video frame (red triangles). E: An enlarged view of the overlapped triangles of the current and next video frame and their incentres. For each triangle the displacement was calculated based on incentre displacement between two consecutive frames.

The local displacement of the skin within the contact area was obtained in a three-stage process: 1) In the first step, the position of valid fingerprint features extracted in sequences of fingertip contact images were segregated (Fig. 2B). 2) Using the valid tracked features of the fingerprint Delaunay triangles were constructed for the consecutive frames (Fig. 2C and 2D). The incentre for each triangle was identified and saved for each frame. Using the triangle incentres, displacement vectors were computed for each triangle within each frame (Fig. 2E). The area for each triangle was then measured. 3) In the third stage, the slipped triangles were identified by using a displacement threshold of one pixel (Barrea et al., 2018). Any vector with displacement greater than one pixel was considered as a slip and the triangle was regarded as a slipped triangle. The displacement vectors of the incentres of the triangles were calculated for each frame.

### Skin displacement magnitude

The skin displacement magnitude was calculated by assigning each pixel within a Delaunay triangle with a displacement vector equal to the movement of the triangle’s incentre. The triangle displacement in the current frame was calculated with respect to its position in the previous frame. We calculated three different skin movements: i) The *total and net displacement* that the fingertip skin moved between the first and last video frames; ii) The *displacement jitter* characterising deviation of trajectory from net displacement vector between the start and the end of the movement and iii) The *divergence*, which in our study denotes skin displacement in each video frame which doesn’t contribute to the net skin displacement within the contact area such as radial inside-out skin displacement.

#### Total and net displacement

First, we calculated the pixel displacement vectors 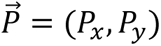 and their norms 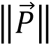 for each pixel *i* in each frame *j*. Thus, each pixel displacement vector represents the vector difference of a coordinates of a point on the skin between two consecutive frames. All the pixel displacement vectors are summed for each frame and represented as *f*_*net_j_*_ and their norms are summed and represented as *f*_*total_j_*_ mathematically:

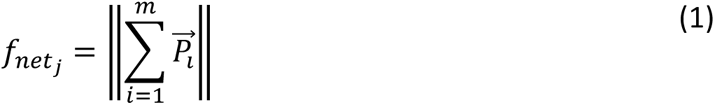

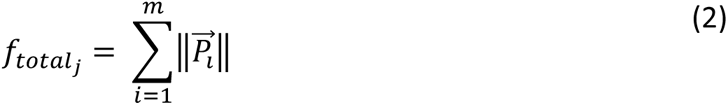

where *m* is the total number of skin contact area pixels in the frame.

#### Displacement jitter

The displacement jitter was quantified by taking a norm of the net displacement vector estimated for each frame then summed across all k frames and subtracting the norm of the net displacement vector between start and end frames estimated as vector sum of all pixel displacements:

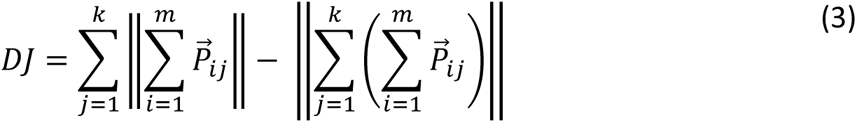

where *k* represents the total number of frames.

#### Divergence

The skin displacement in a divergence pattern, *D*, in our study was calculated by taking the difference between the total cumulative displacement and the net cumulative displacement within each frame and then summed across all the frames analysed:

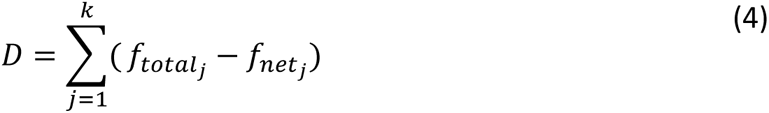

### Statistical analysis

First, we estimated a quotient Q_µs_ of static coefficients of friction µ_s_ for each pair of frictional stimuli. A one-sample t-test was performed to confirm that the Q_µs_ was significantly different from 1 and thus µ_s_ for the two stimuli in a pair are different (where Q_µs_ =1 would indicate that the measured µ_s_ for two stimuli in a pair are equal). Analyses of variance (ANOVAs) and post hoc paired sample tests (with Bonferroni corrected values for multiple comparisons) were performed if comparison of more than two groups was required. Fractional degrees of freedom are reported accordingly to the Greenhouse-Geisser correction when Mauchly’s Test of Sphericity showed that the assumption of sphericity had been violated. When D’Agostino & Pearson normality test (p < 0.05) indicated that the data was not normally distributed, instead of ANOVA the Friedman test was used. Wilcoxon matched-pairs signed rank test was used as a nonparametric alternative to t-test when differences between two groups had to be evaluated.

## Results

Frictional properties of the contact between the fingertip skin and surface are individual and depend upon several variable factors including skin mechanical properties and the amount of sweat. Thus, we first had to assess the actual frictional difference and efficacy of ultrasonic friction modulation between the pairs of stimuli. Then we report subjects’ ability to differentiate between two frictional levels by reporting which surface felt more slippery when reaching and touching the surface (reach-and-touch condition). By means of biomechanical analyses of the fingertip skin strain patterns during the touch, we observe that the source of sensory input determining subjects’ ability to differentiate slipperiness of two surfaces relates to the presence of partial slips. In the follow-up experiment, in which we restrict fingertip movement jitter (supported touch), we determine that natural movement kinematics is at the heart of the friction sensing mechanism at the initial touch.

### Reach-and-touch condition

#### Frictional effect achieved by friction reduction device

The frictional measurements were obtained by subjects contacting the friction reduction device surface and sliding their fingers in the proximal direction. The mean measured coefficient of static friction (µ_s_) during the reach-and-touch experimental block was 0.65 ± 0.27, 0.48 ± 0.22 and 0.29 ± 0.14 (mean ± SD) for high (H), medium (M) and low (L) friction conditions, respectively. The ratio of the larger measured µ_s_ to the smallest measured µ_s_ for the two stimuli in a pair—a quotient Q_µs_— varied amongst subjects, ranging between 1.12 and 1.61 for H vs M, between 1.42 and 2.20 for M vs L, and 1.70 and 3.51 for H vs L friction levels (min-max respectively) (Fig. 3A). A one-sample t-test for each of the frictional combinations indicated that Q_µs_ was significantly different from 1 (H-M, p = 0.0001; M-L, p < 0.0001; H-L, p < 0.0001, n =11) indicating that the µ_s_ of the stimuli presented in every pair were significantly different. It is also apparent that the largest measured frictional differences were in H-L followed by M-L and the smallest differences were measured in H-M pair of stimuli. It should be noted that exact frictional differences cannot be controlled and will inevitably vary between subjects and from trial to trial.

**Fig. 3.**
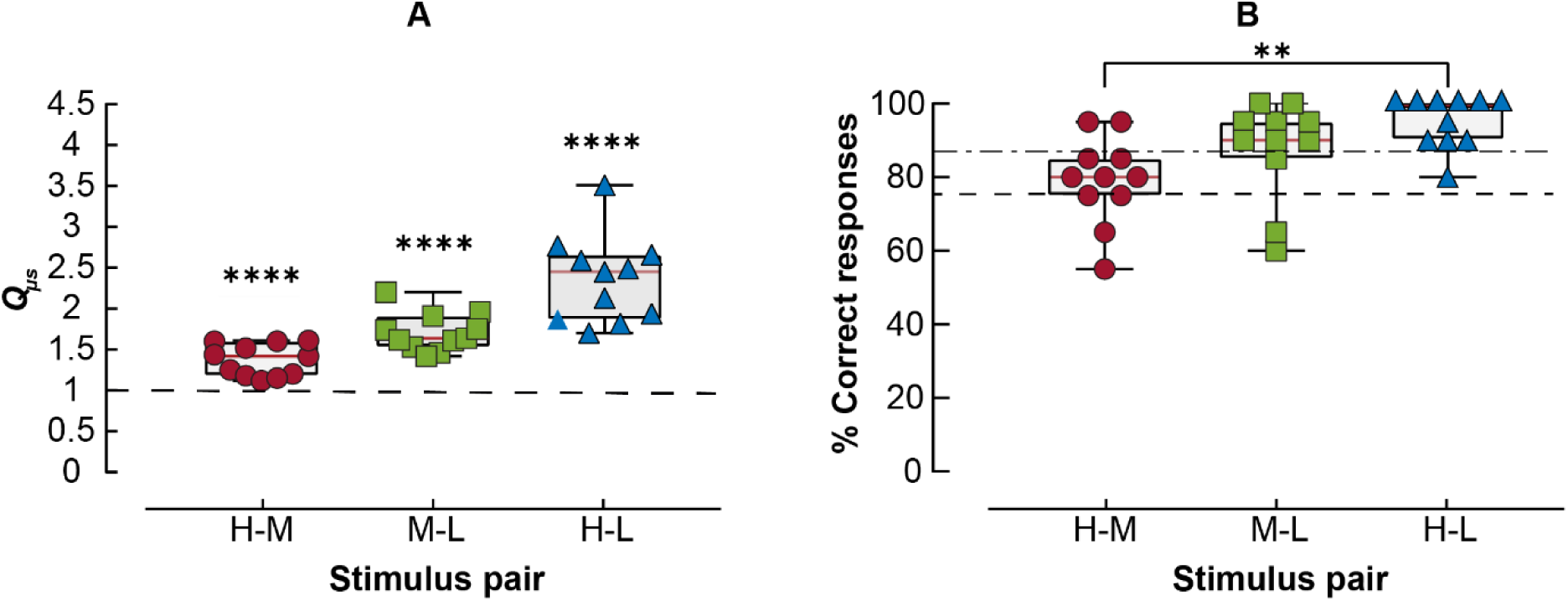
Friction discrimination in reach-and-touch condition. A: Boxplots displaying the median and quartile range of Q_µs_ (quotient of the larger and smaller mean µs for the two stimuli in a pair) measured for individual subjects (n= 11). Dashed horizontal black line across the graph is Q_µs_ = 1, i.e., if two stimuli in a pair would have equal µ_s_. ****Significant difference between the mean Q_µs_ and 1 at p < 0.001; one-sample t-test. B: Boxplots displaying the median and quartile range of percentages of correct responses across friction pairs for reach-and-touch condition. Individual subject data is shown by symbols (n = 11). Dashed horizontal black line represents selected 75% performance threshold level. Dash-dotted line represents the mean of all subjects across stimulus pairs. ** indicates p <= 0.001 (Bonferroni corrected)

#### Subject ability to discriminate friction

The friction discrimination ability of the individual subjects during the reach-and-touch condition for the three pairs of stimuli are shown in Fig. 3B. The overall accuracy averaged across subjects was 87 ± 8% (mean ± SD; n=11). For the pairs of stimuli with the largest frictional difference (H-L) all eleven subjects performed above 75% performance level (95 ± 6%). The discrimination performance between the two stimuli with a smaller frictional difference, H-M and M-L, remained high and was 79 ± 11% and 87 ± 13%, respectively (mean ± SD). There was a significant difference between the performance of the subjects among the three pairs of stimuli (one-way repeated-measures ANOVA, F (1.594, 15.94) = 11.58, p = 0.0013). Post hoc analyses indicated that the performance differed only between stimulus pairs of highest and smallest frictional difference (H-M vs H-L, p = 0.0010) and was similar between pairs of intermediate differences (H-M vs M-L, p = 0.1764; M-L vs H-L, p = 0.0709; Bonferroni-corrected multiple comparisons). Only two subjects S3 and S4 had performance below the 75% threshold (55% and 65%, respectively) in H-M and subjects S3 and S2 in M-L pairs of stimuli (65% and 60%, respectively). The possible reason could be that the frictional difference between the low performing pairs for S3 was relatively small. Q_µs_ measured for the H-M pair on average was 1.25 and Q_µs_ measured for M-L pair was 1.46 which is about the lowest in comparison to other subjects. The performance of these subjects in H-L trials with larger frictional differences was well above 75 percent performance level.

#### Skin deformation leading to partial and full slips

To identify the role of friction-dependent skin deformation and localized slips that signal frictional properties of the surface, we used the fingerprint image analyses and tracked displacement of identified fingerprint image features. Subjects reached and touched the stimulus surface being instructed that sliding lateral finger movements are not allowed and they must touch the surface in a similar manner as when they would grip an object for manipulation. As friction can only be measured when there is a relative motion between the contact surfaces, partial or full slip, first we computed the portion of the fingerprint area that slipped over the time course of making the contact. The point in time when the largest amount of slip occurred potentially being the best source of information for shaping the friction perception. The relative size of the fingertip skin contact area that slipped while touching the surface over the duration of the trial is shown in Fig. 4A.

**Fig. 4.**
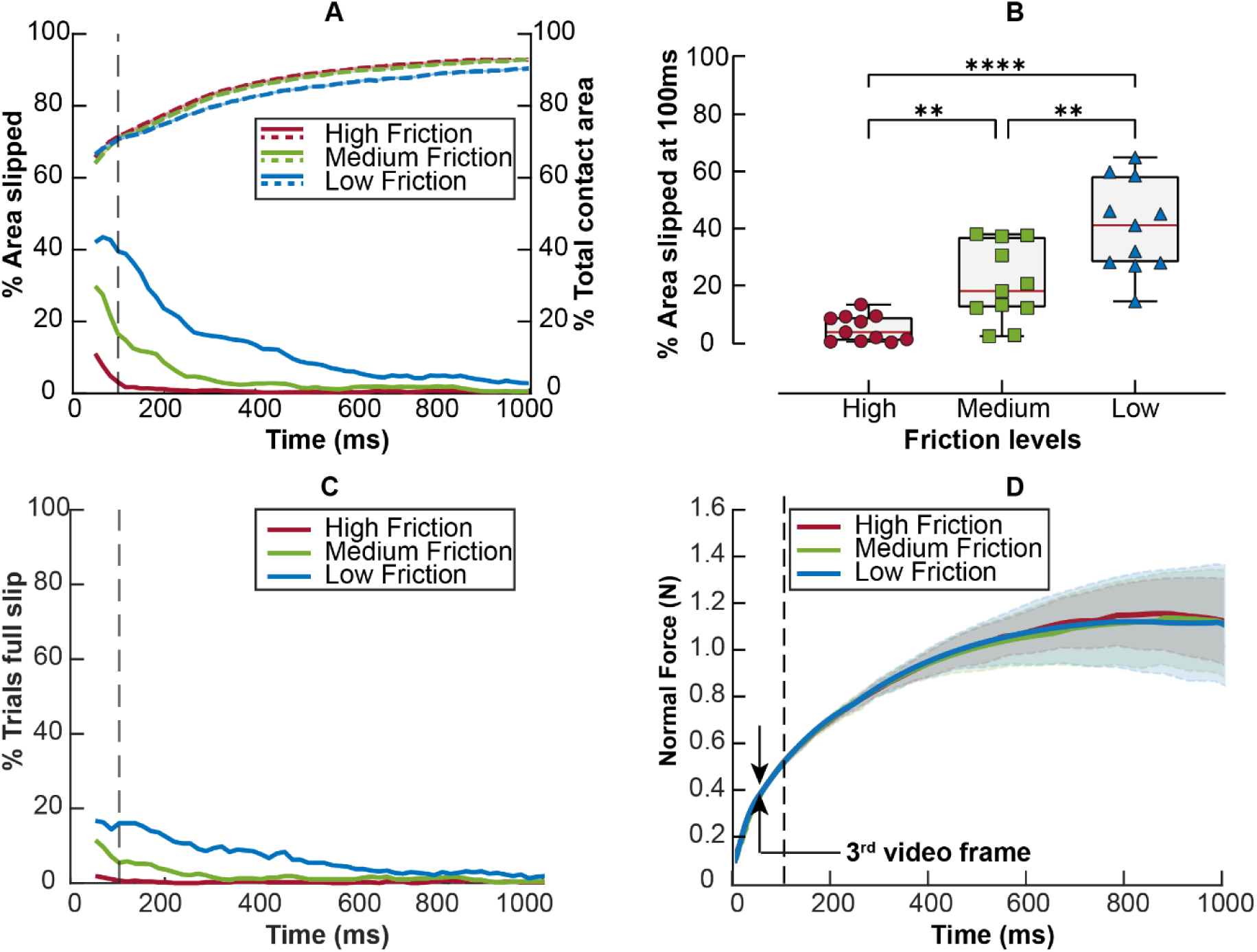
Temporal evolution of fingerprint contact area slipped and normal force development in reach-and-touch condition. A: Mean of the percentage area slipped during the reach-and-touch movement from the initial touch over the duration of the trial (colour-coded lines at the bottom of the graph). The right-side y-axis and dashed lines at the top represent the total contact area growing throughout the touch trial. B: Boxplots displaying the median and quartile range of the percentage area slipping for individual subjects after the first 100 ms of touch, corresponding to the dashed vertical line in panel A (n = 11). **** indicates p <= 0.00001 (Bonferroni corrected). C: Proportion of trials in which full slip was detected at each time point during the trial. D: Normal force development over the duration of contact: The vertical line indicates the 100 ms time point after the initial contact. Solid lines indicate averages across all subjects (n = 11) and shaded areas are standard deviations.

Regardless of frictional condition, 100 ms after touching the surface the area of fingertip skin contacting the surface was 70% of its maximum size reached later during the trial. However, about 40% of the current skin contact area was slipping in the low friction condition, but only 10% was slipping in the high friction condition. This indicates that the fraction of partial slippage relative to the total skin area contacting the surface at this time point reflected the frictional condition. The fingerprint image analysis at the very first frame, when contact is detected, was not reliable because at the very beginning of the touch, while the contact force was very low, the fingerprint features were not discernible yet. Therefore, it cannot be excluded that full slips predominate during this period. The slip detection analyses are performed for feature displacements detected between video frames 2 and 3 (at 60 fps this is 49.98 ms after contact is detected; Fig. 4). The largest fraction of skin area slipping was observed during the first 100 or 200 ms of the contact, which is the time interval most critical for obtaining frictional information to control fingertip forces during object manipulation. The amount of initial slippage was clearly related to the frictional condition: a larger part of the contact area slipped when the friction was low (L) and conversely a lesser amount of skin slipped when the friction was high (H). With the high friction surface, typically all parts of the skin maintained stable contact after the first 100 ms. Fig. 4B summarizes the percentage area slipped and variability between subjects at 100 ms after the contact was made. It can be clearly seen that the size of the slipped portion of the contact area reflected the friction modulation level. After the first 100 ms on average 40.49 ± 16.08%, 20.43 ± 13.54% and 5.02 ± 4.65% of contact area was slipping with H, M and L frictional surfaces, respectively. A one-way repeated-measures ANOVA indicated that the difference was significant (F(1.705, 17.05) = 32.57, p = 0.0001). Post hoc analyses revealed that the area slipped was significantly different between two surfaces presented in pairs H-M, M-L and H-L (H<M, p = 0.0031; M<L, p = 0.0053; H<L, p = 0.0001, respectively; Bonferroni-corrected for multiple comparisons). In the majority of instances, these were partial slips.

The probability of having a full slip (defined as at least 90% of the contact area slipping) is shown in Fig 4C. It is apparent that only 16% of the trials in the low friction condition have full slips at the start of the trial, which shows that subjects have followed the instructions, and this is not an intentional exploratory lateral sliding movement. After this initial period, the contact stability increased, and full slips were rare. The average normal force increase rate during the first 100 ms of contact was 6.21 ± 3.44, 6.24 ± 3.38, 6.92 ± 4.06 N/s (mean ± SD) during H, M and L frictional levels, respectively, thus very similar and not affected by the frictional level (F (1.580, 15.80) = 1.486, p = 0.2532; one-way ANOVA; Fig. 6). At the time point of the third video frame, when the image processing began, the normal force already reached about 0.4 N on average (Fig. 4D).

#### Supported touch

Based on the observations above, we established that, in the absence of intentional exploratory movements, some skin displacement and partial slips were unavoidably present. This indicates that tangential components of the movement kinematics, while being small, could potentially be sufficient to enable subjects to evaluate and discriminate frictional properties of the surfaces. It is supported by our previous study that sub-millimetre range displacement between the object’s surface relative to the rigid nail-phalangeal bone complex may have such an enabling effect. To test whether movement kinematics indeed play a decisive role in friction sensing and perception, we designed an experiment in which we restricted the trajectory of finger movement by introducing a hand support, thus reducing natural, but experimentally unwanted lateral movements. With the introduction of the hand support, no reaching movement was present. It also served to minimize the effects of physiological tremor originating from the large arm muscles. Thus, any tangential finger movements relative to the surface were minimized, constraining it to a straighter path. This condition is termed “supported touch condition”, where the stimulus surface is touched by moving a finger while the palm of the hand and last three fingers enclose the hand support rod (Fig. 1C). We start by reporting the subjects’ performance changes after the introduction of the hand support, then we analyse the extent of changes in skin displacement, and finally to establish causality between skin displacement and subjects’ performance we analyse whether stochastic trial-to-trial variation in displacement can be linked to subjects’ performance.

#### Frictional effect on skin from friction reduction device

The static coefficient of friction measured for each of the three rendered friction levels (H, M and L) was different among subjects. The measured coefficient of friction (µ_s_) was 0.65 ± 0.28, 0.53 ± 0.25 and 0.35 ± 0.18 (mean ± SD) for supported touch. The ratio of the larger measured µ_s_ to the smallest measured µ_s_ (i.e., Q_µs_) for the two stimuli in a pair varied amongst subjects, ranging between 1.05 and 1.53 for H-M, between 1.25 and 2.06 for M-L, and 1.32 and 3.03 for H-L (min-max respectively) during the supported touch condition (Fig. 5A). A one-sample t-test for each of the frictional combinations indicated that there was significant difference between the mean Q_µs_ measured in the supported touch condition (H-M, p = 0.0003; M-L, p < 0.0001; H-L, p < 0.0001, n = 11), meaning that the µ_s_ of the stimuli presented in pairs were significantly different.

**Fig. 5.**
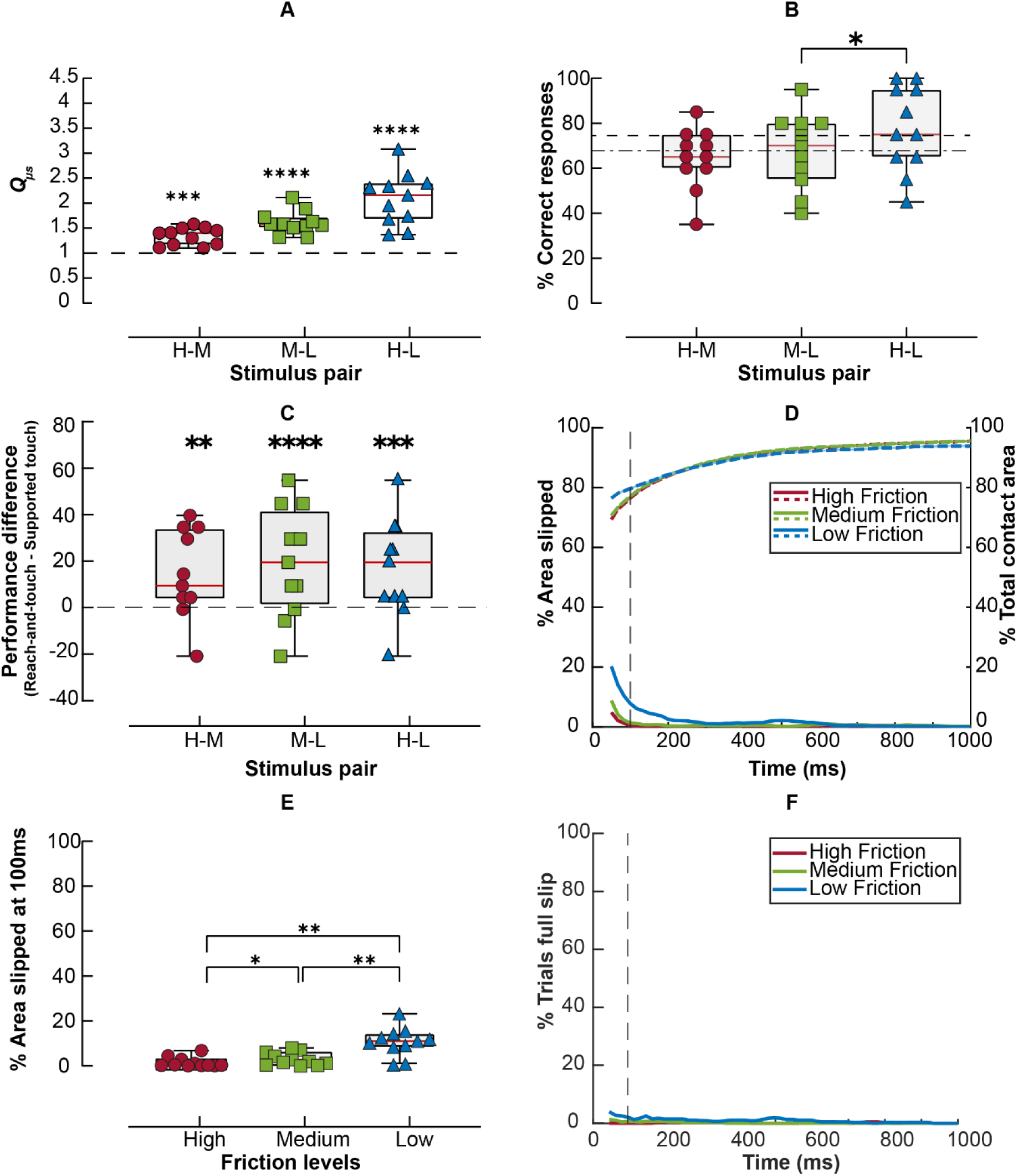
Subjects’ performance and fingerprint contact area slipped in supported touch condition A: Boxplots displaying the median and quartile range of the quotient of the coefficient of friction measured for individual subjects (n= 11). Symbols show data from individual subjects (n = 11). Dashed horizontal black line across the graph is Q_µs_ = 1, i.e., if two stimuli in pair would have equal µ_s_. ****Significant difference between the mean Q_µs_ and 1 at p < 0.001; one-sample t-test. B: Boxplots displaying the median and quartile range of percentages of correct responses across friction pairs for the supported touch condition. Dashed horizontal black line represents 75 percent performance threshold level. Dash-dotted line represents the mean performance of all subjects across all stimulus pairs. C: Boxplots displaying the median and quartile range of performance difference between the reach-and-touch condition and the supported touch condition across friction pairs. **** Indicates p <0.0001 (Bonferroni corrected) D: Mean of the percentage area slipped during the reach-and-touch movement from the initial touch over the duration of the trial (colour-coded lines at the bottom of the graph). The vertical line indicates the initial phase of the contact at 100 ms. The right-side y-axis and dashed lines at the top represent the total contact area growth over the duration of the touch trial. E: Boxplots displaying the median and quartile range of the percentage area slipping for individual subjects after the first 100 ms of touch corresponding to the dashed vertical line in panel A (n = 11). F: Proportion of trials in which full slip was detected as a function of time after the initial contact. The vertical dashed line indicates 100ms after initial contact with the surface.

**Fig. 6.**
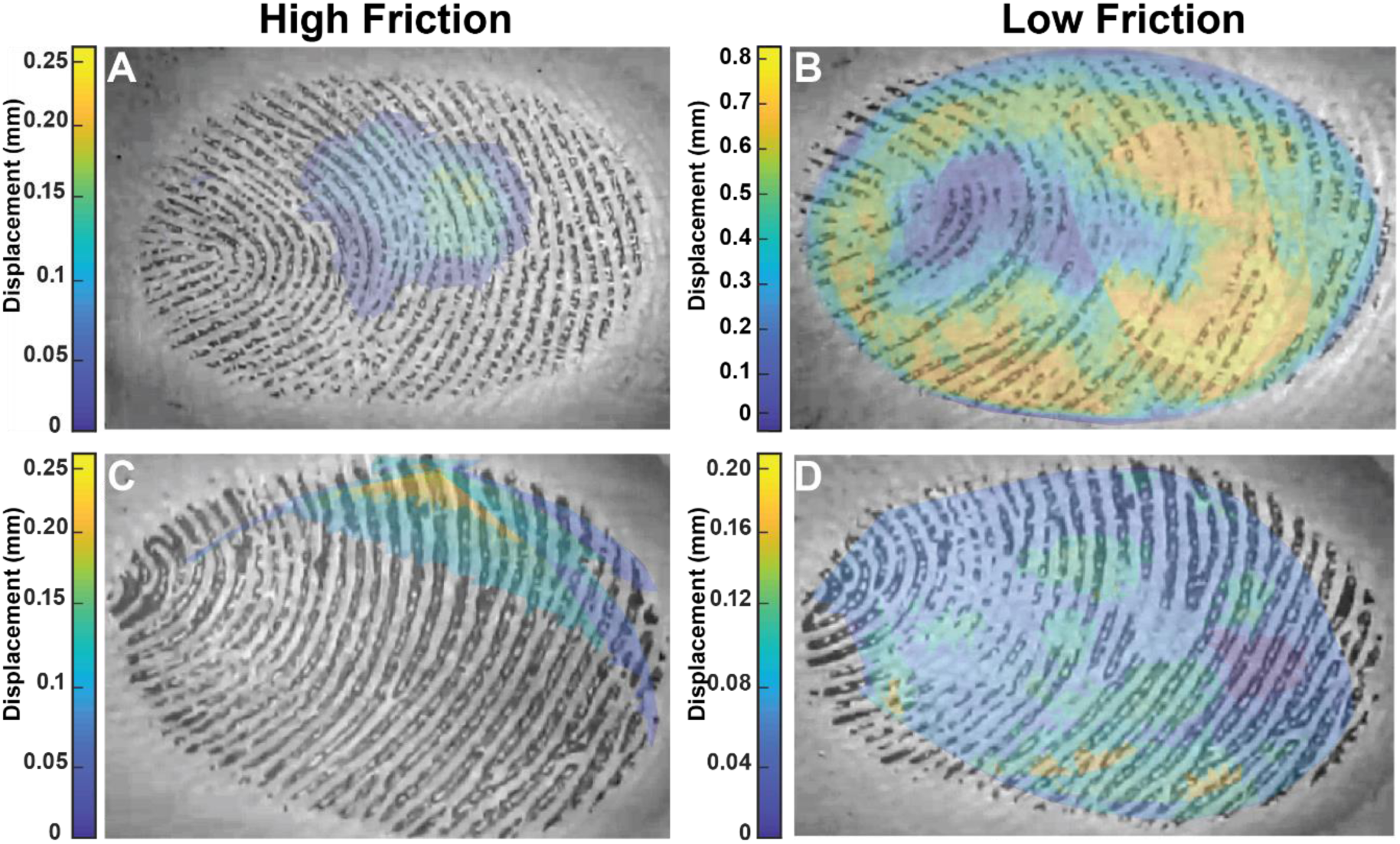
Local displacement magnitude pattern of the fingertip skin touches high and low friction surfaces during reach-and-touch and supported touch condition trials. The heat map represents the local displacement magnitude measured between the first video frame when each fingerprint feature was identified, and the last video frame overlaid on top of fingerprint image. A and B: high friction and low friction in reach-and-touch condition. C and D: high friction and low friction in supported-touch condition.

#### Subjects’ ability to discriminate friction

Due to the introduction of the hand support, the friction discrimination ability of the individual subjects significantly deteriorated (Fig. 5B). For the pairs of stimuli, even with the largest frictional difference (H-L), only five out of 11 subjects performed above the 75 percent threshold. On average, the performance for the H-L friction pair was just above 75% (77.73 ± 18.89%) and considerably lower than in the reach-and-touch condition (95 ± 6%; mean ± SD) for the pairs of stimuli with the largest frictional difference (H-L). The discrimination performance between two stimuli with smaller frictional differences H-M and M-L on average was below the 75 percent threshold, being 67.73 ± 16.64% and 64.55 ± 13.50%, respectively. On average, across three levels of frictional differences, the subjects’ performance was 66.14% ± 12.01%. One-way repeated-measures ANOVA (F (1.535, 15.35) = 4.233, p = 0.0429) and post hoc analyses indicated that the only difference in performance was between the pairs of smallest and largest frictional differences (p = 0.0341 for M-L vs H-L; Bonferroni-corrected multiple comparisons). Two-way-repeated-measures ANOVA (F(1,10) = 9.648, p = 0.011; Fig. 5C) indicated that the performance differed significantly between the two touch conditions. There was no interaction between the touch conditions and the rendered friction levels (F(2,20) = 0.538, p = 0.591). The poor performance of subjects under constrained movement kinematics was explained by a significantly reduced amount of typically submillimeter range lateral movements and by the extent of partial slips.

#### Skin deformation leading to partial and full slips

The constraints imposed on the finger movement kinematics substantially affected skin displacement magnitude after contact with the surface, which might explain the poor performance of subjects in discriminating the friction difference of the stimulus pairs. The mean percentage of area slipped for all the subjects over the 1 second duration of the contact is shown in Fig. 5D. Overall the contact with the stimulus surface was very stable, with reduced movement of the fingers over the duration of the whole trial, as compared to the unrestrained reach-and-touch condition. It can be observed that after the initial touch with a surface, there seems to be no further slip or even partial slip present in the supported touch condition, as compared to the reach-and-touch condition where small amounts of partial slip were present throughout the trial. At 100ms after initial contact was made (Fig. 5E) on average 10.70% ± 6.35%, 3.08% ± 2.87%, and 1.55% ± 2.24% of the contact area was slipping during the H, M, and L frictional levels, respectively. One-way repeated-measures ANOVA indicated that the mean percent area slipped was significantly different between H, M, and L frictional levels at 100ms after the initial contact was detected (F(1.066,10.66) = 21.89, p = 0.0006). Post hoc analyses revealed that the area slipped was significantly different between any two rendered friction levels (H<M, p = 0.0316; M<L, p = 0.0024; H<L, p = 0.0025, respectively; Bonferroni-corrected for multiple comparisons). Regardless of frictional effects, the size of the area that slipped was relatively small and appeared to be insufficient for the subjects to base the decision on which surface was more slippery. In comparison to the reach-and-touch condition, the area slipped in supported touch was only 26%, 15% and 30% from what it was in the reach-and-touch condition. Full slip was observed only with the low friction surface in 5% of trials (Fig. 5F). Full slips were almost fully absent with the medium and high friction rendered surfaces. The average normal force increase rate during the first 100ms of contact was 4.26 ± 3.18, 4.09 ± 3.11, 4.70 ± 3.30 N/s during H, M, and L frictional level, respectively, thus very similar and not affected by the frictional level (F (1.724, 17.24) = 1.795, p = 0.1980; One-way ANOVA).

#### Effect of hand support on skin displacement magnitude and slip patterns

To further demonstrate how fingertip skin displacement and slip patterns reflect frictional differences, fingerprint image feature tracking was performed. Fig. 6 shows example patterns of local skin displacement magnitude when touching surfaces with high and low friction (panels A vs B; C vs D) and compares the effects between reach-and-touch vs supported touch conditions (A, B vs C, D panels). When touching the high friction surface, the local skin displacement is negligible and does not exceed a magnitude of about 0.1 mm. However, with the low friction surface, at some time during the trial, most of the skin area in contact with surface has slipped; although, the extent of slip displacement depended on the type of touch. During the reach-and-touch condition, the displacement is substantial, reaching about 0.8-1.0 mm in magnitude, while in supported touch the slip is considerably smaller about 0.1mm.

#### Skin displacement measured over the duration of whole trial

The total displacement of skin segments over the whole duration of the trial was much less for the supported touch condition in comparison to the reach-and-touch condition. The relative fraction of the total skin displacement in the supported touch condition was 0.19 (0.11 – 0.45), 0.08 (0.04-0.32) and 0.09 (0.06-0.18) (medians, quartiles) from what it was in the reach-and-touch condition, with H, M and L friction levels, respectively (Fig. 7A). As an exception, in two out of eleven subjects the movement resulted in larger skin displacement in the support condition. Wilcoxon matched-pairs signed-rank test performed on the measured values of the total displacement of skin segments between reach-and-touch condition and supported touch condition indicated that statistically there was a significant difference when compared at each of three friction levels (H, p = 0.0419; M, p = 0.0244; L, p =0.0009).

**Fig. 7.**
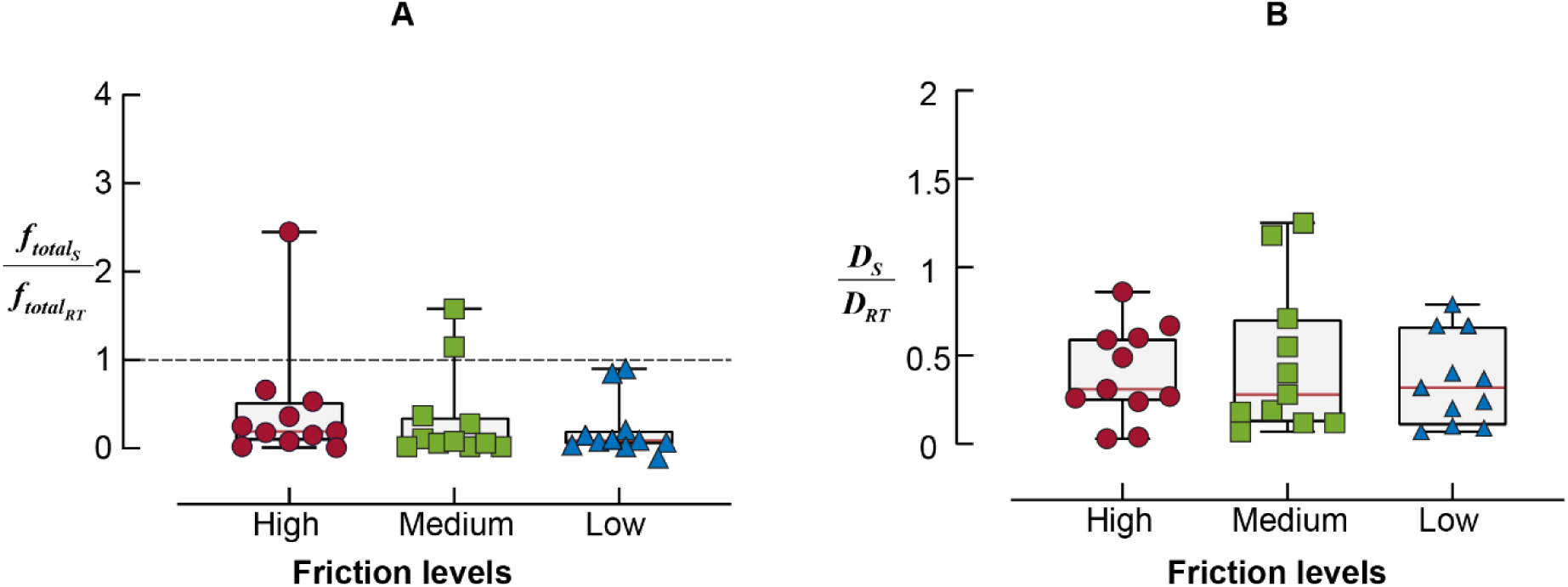
Comparative skin displacement analysis between two touch conditions: total skin displacements and skin divergence. A: Boxplots represent the median and quartile range of the fraction of the total skin displacement after the hand support was introduced for each of three friction levels. *f*_*total_S_*_ represents the total skin displacement measured during the supported touch condition and *f*_*total_RT_*_ represents the total skin displacement measured during the reach-and-touch condition. B: Boxplots represent the median and quartile range of the fraction of skin divergence after the hand support was introduced for each of three friction levels. *D*_*S*_ represents the divergence measured during the supported touch condition and *D*_*RT*_ represents the divergence measured during the reach-and-touch condition.

In the local skin displacement pattern, we specifically discerned a divergence and small displacement jitter, which might partly arise from physiological tremor. Radial divergence of the skin will occur during the skin-object contact because of the approximately hemispherical shape of the fingertip. The extent of divergence depends on the friction. The fraction of divergence relative to the total displacement during the reach-and-touch condition was 0.15 (0.11 - 0.30), 0.07 (0.03 - 0.12), and 0.03 (0.02 - 0.07) (medians, quartiles) during H, M and L friction levels, respectively. In the supported touch condition, the divergence measured relative to the total skin displacement was 0.25 (0.14 - 0.30), 0.07 (0.03 - 0.12), and 0.03 (0.02 - 0.07) (medians, quartiles) during H, M and L friction levels, respectively. In both grip configurations, the effect of friction on divergence was statistically significant (Q = 18.73, p < 0.0001 reach-and-touch condition; Q = 11.64, p = 0.0014 supported touch condition; Friedman test). The relative difference of the divergence between the two conditions (i.e., the ratio of the divergence measured for the supported touch condition to the divergence measured for the reach-and-touch condition) was 0.31 (0.25 - 0.60 quartiles), 0.28 (0.15 – 0.63), and 0.32 (0.15 – 0.53) (medians, quartiles) during H, M and L friction levels, respectively (Fig. 7B), indicating that divergence observed in the supported touch condition was about one-third of what it was in the reach-and-touch condition. This indicates that increased lateral movement present in the reach-and-touch condition facilitated the release of radial stress within the contact area.

The fractional contribution of displacement jitter of the fingers to the total skin displacement was 0.08 (0.06 – 0.14), 0.11 (0.08 – 0.14) and 0.17 (0.11 – 0.27) in reach-and-touch condition and 0.09 (0.03 – 0.13), 0.08 (0.04 – 0.17) and 0.11 (0.09 – 0.16) in the supported touch condition (medians, quartiles with H, M and L friction levels). This indicates that, overall, the contribution of the displacement jitter to the total skin displacement in most cases was negligible.

#### Subjects’ ability to discriminate frictional properties is correlated with stochastic variation of skin area slipped

To evaluate whether subjects’ performance was influenced by the stochastic variation in the size of the area slipped we analysed and compared area slipped between two surfaces in the pair during trials in which subjects correctly and incorrectly identified the more slippery surface. First, we estimated Difference Percent Slipped area by subtracting the area slipped (expressed as percentage of the total area) when touching the less slippery surface from the area slipped for the more slippery surface in the pair. Data from both reach-and-touch and supported touch conditions were used in these analyses. The difference in Percent Slipped area for each pair of stimuli during correct and incorrect trials is shown in Fig. 8, left panels. It can be observed that when the responses were correct, the Difference Percent Slipped area was consistently higher in the trials where subjects identified frictional difference correctly as compared to the trials where subjects were unable to discriminate frictional difference (Fig. 8, right panels). This demonstrates that the subject’s ability to discriminate friction of two surfaces was clearly related to and likely to be determined by the stochastic variation in the size of partial slip differences.

**Fig. 8.**
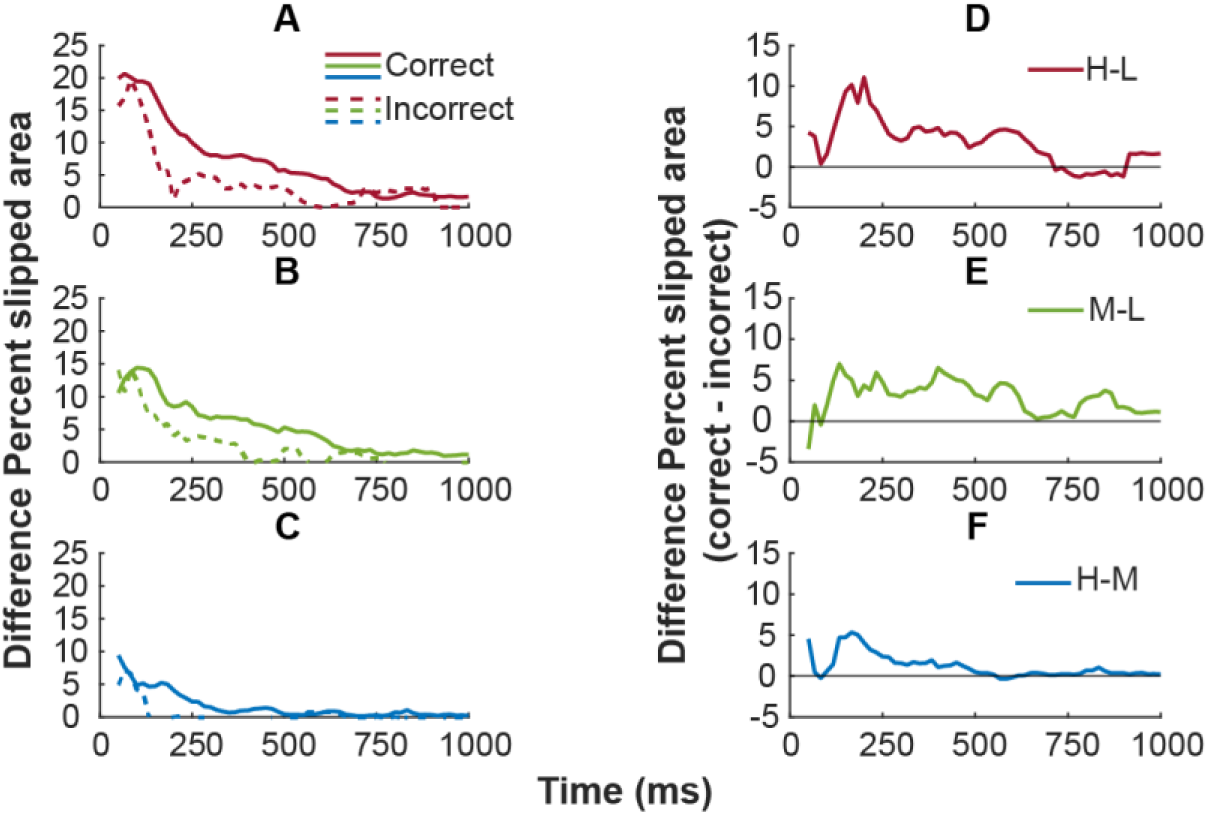
Difference Percent Slipped area between two surfaces in the friction pair over the course of trial during correct and incorrect trials. Difference Percent Slipped area for (A) H-L, (B) M-L, and (C) H-M frictional pairs. Solid lines represent correct trials and dashed lines incorrect trials. The panels on the right (D-F) show the Difference Percent Slipped area in incorrect trials subtracted from the correct trials.

## Discussion

Sensing friction between fingertip skin and a surface may serve two purposes—to explore characteristics of a material or to control grip forces when manipulating an object held in the hand. The same physical feature, friction, is sensed under different sensory conditions. To evaluate the physical characteristics of a material, we slide our fingers over the surface creating a perceptual experience. Object manipulation, however, does not permit such exploratory movements and frictional information should be obtained just by touching the surface. Grip force adjustments to friction are largely automatic (Birznieks et al., 1998) and thus in principle do not necessitate cognitive involvement. Nevertheless, some associated awareness of grip surface slipperiness is present, which is perceptually similar to that felt during exploratory sliding or rubbing finger movements over surfaces. Knowledge of slipperiness can be critical for the selection of a safe and achievable action plan ensuring that while performing an intended manipulation the required grip forces to hold an object or tool safely would not exceed the hand’s physical ability to produce them or not exceed the breakage point of fragile objects.

Previous studies have demonstrated that humans’ ability to differentiate the frictional properties of two otherwise identical materials when touching them might be challenging and depend on multiple factors. In the case of well controlled stimuli—when robotic manipulator brought a surface in contact with the fingertip skin—the subjects were unable to differentiate them based on frictional their properties. This was demonstrated when friction was changed using a friction modulation device (Wiertlewski et al., 2016) with either smooth surfaces or with surface textures (Khamis et al., 2021b). Even changing the approach angle of the robotic manipulator from perpendicular to 20° and 30° from normal (producing 8.9° and 19° force angles and 0.47 ±0.07 mm lateral movement after the first contact was detected) didn’t improve performance. In the follow up study using the same experimental setup in which a robotic manipulator brought a surface to an immobilised finger it was found that a submillimeter range lateral movement as small as 0.5 mm and, in some subjects, just 0.2 mm of the surface relative to the affixed rigid nail-phalangeal bone complex was sufficient to enable the subjects to differentiate surfaces based on their slipperiness (Afzal et al., 2022). Such small tangential movement deviations from a straight path when making contact with a surface are expected to occur during natural movement when humans reach and touch object surfaces. Based on that, in the current study, we tested hypotheses that natural hand movement kinematics would have an enabling effect to sense the slipperiness of smooth surfaces when no other sensory cues are available. It has to be recognised that the fact of the presence of such tangential submillimeter range movements doesn’t mean that it would necessarily enable subjects to differentiate friction due to the stochastic unpredictable nature of these movements in comparison to the experimental setup where applied displacements were reproducible, controlled by a robot, and thus identical. When subjects themselves touch surfaces there would be trial-to-trial variability in several contact parameters, including the magnitude of tangential displacement, adding complexity to the sensory signal from which differences in slipperiness have to be derived.

Our experiments demonstrated that active movements enabled subjects to perceive differences in slipperiness of surfaces which they were previously unable to differentiate in a passive condition (Khamis et al., 2021b). We also demonstrated that in most cases partial slips were sufficient to make the judgement. Full slips occurred in the minority of trials and skin displacements were over very small distances confirming that subjects followed the instructions and didn’t attempt to use exploratory sliding movements. Partial slips were especially prominent in the early phase when contact with the surface was made, thus being the most informative period for receptors to signal frictional properties of the surface. The fingerprint image analyses to identify skin deformation and slips became available only after the first two video frames (∼32 ms) of the contact when contact with a sufficient grip force was made and fingerprint features became discernible. Thus, some finger sliding over the surface might have occurred which potentially could provide rich sensory information. Single afferent recordings in humans (microneurography studies) and psychophysics experiments employing controlled stimuli may reveal the relative importance of these mechanical events.

The subjects’ ability to evaluate slipperiness of surfaces dramatically deteriorated when movement constraints were introduced, and the contact was made by finger movements with limited lateral deviation from a straight path. For most subjects, performance fell to the chance level demonstrating that natural movement kinematics played a decisive role in enabling the subjects to evaluate surface frictional properties. Constrained finger movement paths significantly reduced skin displacement: the size and extent of the partial slips. Furthermore, the analyses of stochastic trial-by-trial variation in skin displacement further confirmed the link between the differences in the size of the partial slip area and subjects’ judgement of differences in slipperiness.

The largest fraction of the skin slipping over the surface was comprised of net unilinear displacement. Other patterns of skin displacement such as the displacement jitter and divergence comprised a minor fraction. The magnitude of skin displacement associated with the divergence was dependent on friction. Importantly, with the natural reach and touch movement the constraints applied to the hand movement kinematics significantly reduced the divergence to about one third of what it was with the natural reach and touch movement.

Thus, natural movement kinematics profoundly enhanced the availability of friction-related sensory cues related to the divergence pattern. This is because the lateral movement promoting partial slips facilitated the release of stress in radial patterns as predicted by the skin mechanical modelling (Willemet et al., 2021). As the magnitude of unilinear skin displacement concurrently caused profoundly larger divergence magnitudes, the relative significance of the two in signalling friction currently cannot be separated. It could be speculated that the ratio between the unilateral displacement and divergence might be exploited to reduce ambiguity in extracting frictional information from the amount of skin displacement, due to the stochastic size of tangential movement. For example, when a relatively larger net skin displacement occurs on a less slippery surface, caused by an overall larger lateral movement than with a more slippery surface. One of alternative possibilities to reduce ambiguity due to variability of contact kinematics in real life situation is that the magnitude of skin displacement must be viewed in the context of the skin stretch pattern caused by lateral movement or torsion in the finger pad (Khamis et al., 2015; Dunilac et al., 2023; Loutit et al., 2023). As in virtual environments, tangential skin stretches within the contact area ranging between 0.25 - 0.75 mm leads to friction perception (Santello et al., 2002; Provancher and Sylvester, 2009; Kamikawa and Okamura, 2018; Suchoski et al., 2018).

The current study has identified the neurophysiological role of active movement, contact kinematics, and slips in how frictional information is obtained and used by the brain for perception. This shows for the first time that small lateral movements as a part of natural motor control during grasping helps inform our perception of friction between the finger pad skin and the object being touched. Our findings can inform engineers of how programmed movement jitter may improve the sensitivity of tactile sensors (Khamis et al., 2019; Huloux et al., 2021; Khamis et al., 2021a) which could measure friction (Chen et al., 2018; Khamis et al., 2018; Ulloa et al., 2022) and in designing tactile displays which make use of friction modulation principles (Gueorguiev et al., 2017; Vardar et al., 2017). This study also has implications in the design of neural prostheses and extends the understanding of tactile sensorimotor control strategies required for dexterous object manipulation (Suresh et al., 2020; Bensmaia et al., 2023).

## Acknowledgments

This project was funded by the Australian Research Council Discovery Grants DP230100048 and DP200100630. We thank Mr. Hilary Carter (NeuRA) for assistance with mechanical works.

